# Essentiality of LD-Transpeptidation in *Agrobacterium tumefaciens*

**DOI:** 10.1101/2024.06.21.600065

**Authors:** Alena Aliashkevich, Thomas Guest, Laura Alvarez, Michael C. Gilmore, Jennifer Amstutz, André Mateus, Bastian Schiffthaler, Iñigo Ruiz, Athanasios Typas, Mikhail M. Savitski, Pamela J. B. Brown, Felipe Cava

**Affiliations:** Department of Molecular Biology and Laboratory for Molecular Infection Medicine Sweden, Umeå Centre for Microbial Research, SciLifeLab, Umeå University, Umeå, Sweden; Division of Biological Sciences, University of Missouri-Columbia, Columbia, Missouri, USA; Genome Biology Unit, European Molecular Biology Laboratory, Heidelberg, Germany

**Keywords:** LD-transpeptidase, LD-crosslink, *Agrobacterium tumefaciens*, peptidoglycan, cell division, Hyphomicrobiales, polar growth

## Abstract

Peptidoglycan (PG), a mesh-like structure which is the primary component of the bacterial cell wall, is crucial to maintain cell integrity and shape. While most bacteria rely on penicillin binding proteins (PBPs) for crosslinking, some species employ LD-transpeptidases (LDTs). Unlike PBPs, the essentiality and biological functions of LDTs remain largely unclear. The Hyphomicrobiales order of the Alphaproteobacteria, known for their polar growth, have PG which is unusually rich in LD-crosslinks, suggesting that LDTs may play a more significant role in PG synthesis in these bacteria. Here, we investigated LDTs in the plant pathogen *Agrobacterium tumefaciens* and found that LD-transpeptidation, resulting from at least one of 14 putative LDTs present in this bacterium, is essential for its survival. Notably, a mutant lacking a distinctive group of 7 LDTs which are broadly conserved among the Hyphomicrobiales exhibited reduced LD-crosslinking and tethering of PG to outer membrane β-barrel proteins. Consequently, this mutant suffered severe fitness loss and cell shape rounding, underscoring the critical role played by these Hyphomicrobiales-specific LDTs in maintaining cell wall integrity and promoting elongation. Tn-sequencing screens further revealed non-redundant functions for *A. tumefaciens* LDTs. Specifically, Hyphomicrobiales-specific LDTs exhibited synthetic genetic interactions with division and cell cycle proteins, and a single LDT from another group. Additionally, our findings demonstrate that strains lacking all LDTs except one displayed distinctive phenotypic profiles and genetic interactions. Collectively, our work emphasizes the critical role of LD-crosslinking in *A. tumefaciens* cell wall integrity and growth and provides insights into the functional specialization of these crosslinking activities.

## Introduction

Most bacteria are surrounded by an essential protective mesh-like structure called the peptidoglycan (PG) or murein sacculus, comprised of glycan chains of repeating β-1,4-linked N-acetylglucosamine (NAG) and N-acetylmuramic acid (NAM) sugars, tethered by peptide crosslinks formed between adjacent peptide side chains attached to NAM.

During growth, expansion of the sacculus requires the coordinated action of synthetic and degradative enzymes that catalyze the insertion of new material into the pre-existing structure. The paradigm in rod-shaped bacteria has been that two protein assemblies target PG biosynthesis at specific times and locations: the elongasome complex inserts new PG along the lateral sidewall whereas the divisome operates at mid-cell to enable cell division (1). The canonical machineries for elongation and division utilize similar protein components, suggesting a shared evolutionary history (2). Specifically, they involve SEDS (Shape, Elongation, Division, and Sporulation) proteins, such as RodA or FtsW, which possess glycosyltransferase activity for extending PG glycan strands, and monofunctional or class B penicillin-binding proteins (bPBP) with DD-transpeptidase activity, such as PBP2 or PBP3, to crosslink peptides in adjacent glycan strands (3–5). Independently from these complexes, PG biosynthesis is further supported by bifunctional class A PBPs (aPBPs), enzymes that have both glycosyltransferase activity and DD-transpeptidase activity (6). In addition to the DD-transpeptidases, many bacteria encode alternative crosslinking enzymes known as LD-transpeptidases or LDTs (7) that do not share sequence homology with PBPs. They present a YkuD-like domain (PFAM 03734) that includes a cysteine as the catalytic nucleophile instead of the conserved serine in PBPs (7). While PBPs form DD or 4,3 crosslinks between their 4th and 3rd amino acids, D-alanine and meso-diaminopimelic acid (mDAP) in Gram-negatives, LDTs catalyze the LD or 3,3 type between the L and D chiral centres of two mDAP residues and can crosslink outer membrane proteins to the PG (Fig. 1A) (8–11). Interestingly, *A. tumefaciens* does not have any homologues to the new family of VanW-domain containing LDTs found in *Clostridioides* (12).

**Fig 1.**
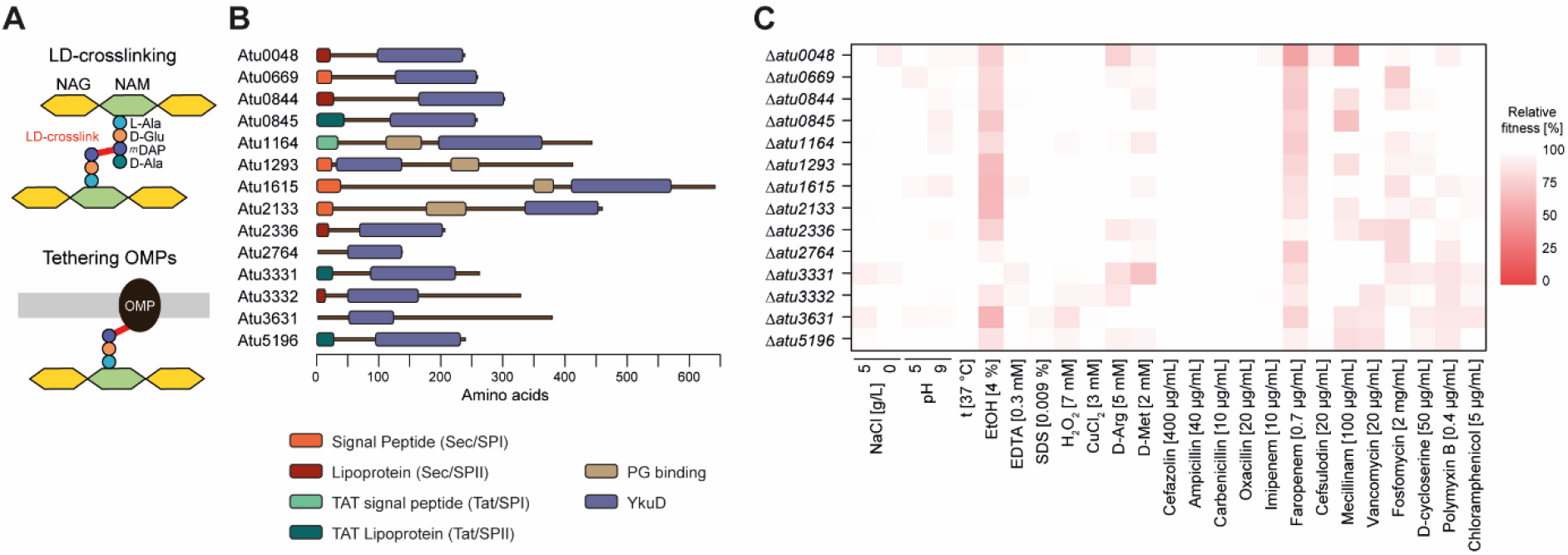
Functional role and contribution of L-D transpeptidases to *A. tumefaciens* fitness. **(A)** Cartoon depicting the major functions of LDTs; formation of LD crosslinks between PG chains and the tethering of outer membrane β-barrel proteins to the PG. **(B)** Predicted signal peptides and protein domain architecture of the 14 putative LDTs in *A. tumefaciens*. **(C)** Heatmap depicting the relative fitness of LDT deletion mutants (compared to the wild type) assessed during growth across a panel of conditions.

In most rod-shaped Gram-negative model bacteria like *Escherichia coli,* the SEDS proteins and monofunctional DD-transpeptidase are essential component of PG biogenesis while LD-transpeptidation is dispensable albeit important for a number of processes such as chemical modification of PG with non-canonical D-amino acids (NCDAA) (8), tethering of outer membrane proteins to the PG (9, 13), toxin secretion (14), lipopolysaccharide translocation (15) and antibiotic resistance (16). However, the absence of the core elongasome components in most polarly growing rods belonging to Actinobacteria and Hyphomicrobiales (aka, Rhizobiales) (17, 18) highlights this pervasive model for elongation is not universal. For instance, this is the case for the plant pathogen *Agrobacterium tumefaciens,* which lacks RodA and PBP2 proteins and instead depends on PBP1A for unipolar growth (19). Moreover, in comparison to *E. coli*, the PG of *A. tumefaciens* is both highly crosslinked and enriched for LD crosslinks (17). LD crosslinks account for ∼1-5 % of cross links in the PG of *E. coli,* while their proportion in *A. tumefaciens* is roughly 30 % (17, 20). The genome of *A. tumefaciens* encodes 14 putative LDTs of which seven are specific to polar-growing species of the Hyphomicrobiales (21). Furthermore, a subset of *A. tumefaciens* LDTs localise to the growth pole during elongation which suggests they could play a role in polar growth (21). However, little is known about the role LDTs play in cell wall homeostasis and polar growth.

We investigated the potential role of LDTs as main contributors to polar PG biosynthesis in *A. tumefaciens* in contrast with the ancillary PG remodelling functions often attributed to these enzymes in other species (7). Here, we show that *A. tumefaciens’* LD-transpeptidases are only partially redundant and inactivation of all of them is lethal. To the best of our knowledge, this is the first reported case of a Gram-negative bacterium for which LD-transpeptidation is essential for survival. We further found that the Hyphomicrobiales-specific LDTs are genetically linked to canonical division proteins and vital for maintaining cell wall integrity and cell shape. Overall, our observations indicate that LDTs are important for polar growth and resistance to cell envelope stress in *A. tumefaciens*.

## Results

### Structural diversity and conservation of *A. tumefaciens* LDTs

The 14 putative LDTs encoded in the *A. tumefaciens* genome feature a conserved YkuD-domain, but their size, predicted structure and the presence of additional signal and attachment domains varies considerably (Fig. 1B, S1). For instance, while two proteins (Atu2764 and Atu3631) lack a predicted signal peptide, the others have some variations of one (i.e., Sec, TAT, and lipoprotein signal peptides), indicating different mechanisms of membrane anchoring and translocation. Additionally, the YkuD-domain can be situated either at the N-terminus or the C-terminus. To ascertain their degree of conservation we compared LDT homologues amongst approx. 50 Pseudomonadota species with putative LDTs numbering between 0 (e.g., *Comamonas testosteroni*) and 21 (*Bradyrhizobium diazoefficiens*) (Fig. S2). Interestingly, among the 6 YkuD-containing proteins (LdtA-F) documented in *E. coli* (15), only LdtD and the endopeptidase LdtF have counterparts in *A. tumefaciens*, namely Atu1615 and Atu1164 for LdtD, and Atu3631 and Atu3332 for LdtF (Fig. S2AB). While some LDTs, such as those mentioned, are widely conserved across Pseudomonadota, seven are predominantly confined to the Hyphomicrobiales and some Rhodobacterales (21). Furthermore, a few lack homologs among the species analyzed (Fig. S2C), implying a distinct evolutionary lineage for these proteins.

### Functional redundancy of LDTs in *A. tumefaciens* is only partial

To investigate the essentiality and function of these proteins, we constructed deletion mutants for each of the 14 LDT genes. None of these individual mutants had any significant defects in growth, morphology, or LD crosslinking (Fig. S3) under standard growth conditions (LB medium + 0.5% NaCl (LB5), 30 °C, aerobic growth), supporting the presumed redundancy of LDTs in *A. tumefaciens*. To further assess their individual contribution to bacterial fitness we subjected these mutants to a panel of diverse physicochemical challenges and antibiotics that challenge the integrity of the bacterial cell envelope (Fig. 1C, Supplementary Table S5, Fig. S3F, Supplementary File 1). In general, the LDT mutants grew as well as the wild type and were largely unaffected. The mutant in *atu0048*, encoding a Hyphomicrobiales- and Rhodobacterales-specific LDT, was the most susceptible across the whole panel of growth conditions. The majority of the LDT mutants were more susceptible to faropenem, a carbapenem antibiotic that decreases the abundance of both LD and DD crosslinks (19). Only the mutant in *atu2336* (the closest homolog to Atu0048, 50% identity) was unperturbed in the presence of faropenem. However, this mutant was more susceptible to fosfomycin, an antibiotic that targets precursor synthesis, whereas Δ*atu0048* was unaffected. Similarly, the LDT mutant strain Δ*atu3331* was insensitive to challenge with 4% EtOH but exhibited the most pronounced response among all 14 mutants to D-methionine, a non-canonical D-amino acid that is synthetically lethal in combination with defects in cell wall biosynthesis genes (22). Collectively, the screen shows that despite a high level of redundancy the LDTs show certain functional specialization.

### Hyphomicrobiales-specific LDTs play major roles in shape determination and cell wall integrity

Given the substantial redundancy observed among the individual LDTs, we clustered them based on their protein sequence similarity, which led to the identification of three distinct groups (Fig. 2A). Group 1 consists of six LDTs, each exhibiting low identity with one another and differing levels of conservation within the Pseudomonadota. Group 2 consists of a single evolutionarily distinct LDT, Atu2133. Finally, group 3 includes the seven LDTs exclusive to Hyphomicrobiales and certain Rhodobacterales. To evaluate the impact of the different groups of LDTs on *A. tumefaciens* fitness and cell wall integrity, we constructed mutant strains in which all the LDTs in each group were deleted. Deletion of group 1 LDT genes (Δgr1) did not alter growth when the bacteria were grown in standard (LB5) or hypoosmotic (LB without added NaCl, LB0) growth conditions (Fig. 2B). Conversely, growth of the Δgr3 strain was severely impaired, especially under hypoosmotic stress. To further distinguish the contributions of the LDT groups to bacterial survival we subjected the group mutants to the same panel of stresses used earlier (Fig. 2C, Supplementary File 1). Growth of Δgr1 mutant was mostly unaffected, though there was a small decrease in its relative fitness compared to wild type for many of the conditions tested. Across the panel of conditions, the fitness of the Δgr3 mutant was severely impaired; in particular, this mutant is highly susceptible to alkaline pH, the presence of D-arginine and many cell wall targeting antibiotics such as cefazolin, ampicillin and carbenicillin. Collectively, these results indicate that the maintenance of cell shape and cell wall homeostasis in *A. tumefaciens* depends more on group 3 LDTs than on group 1 LDTs.

**Fig. 2.**
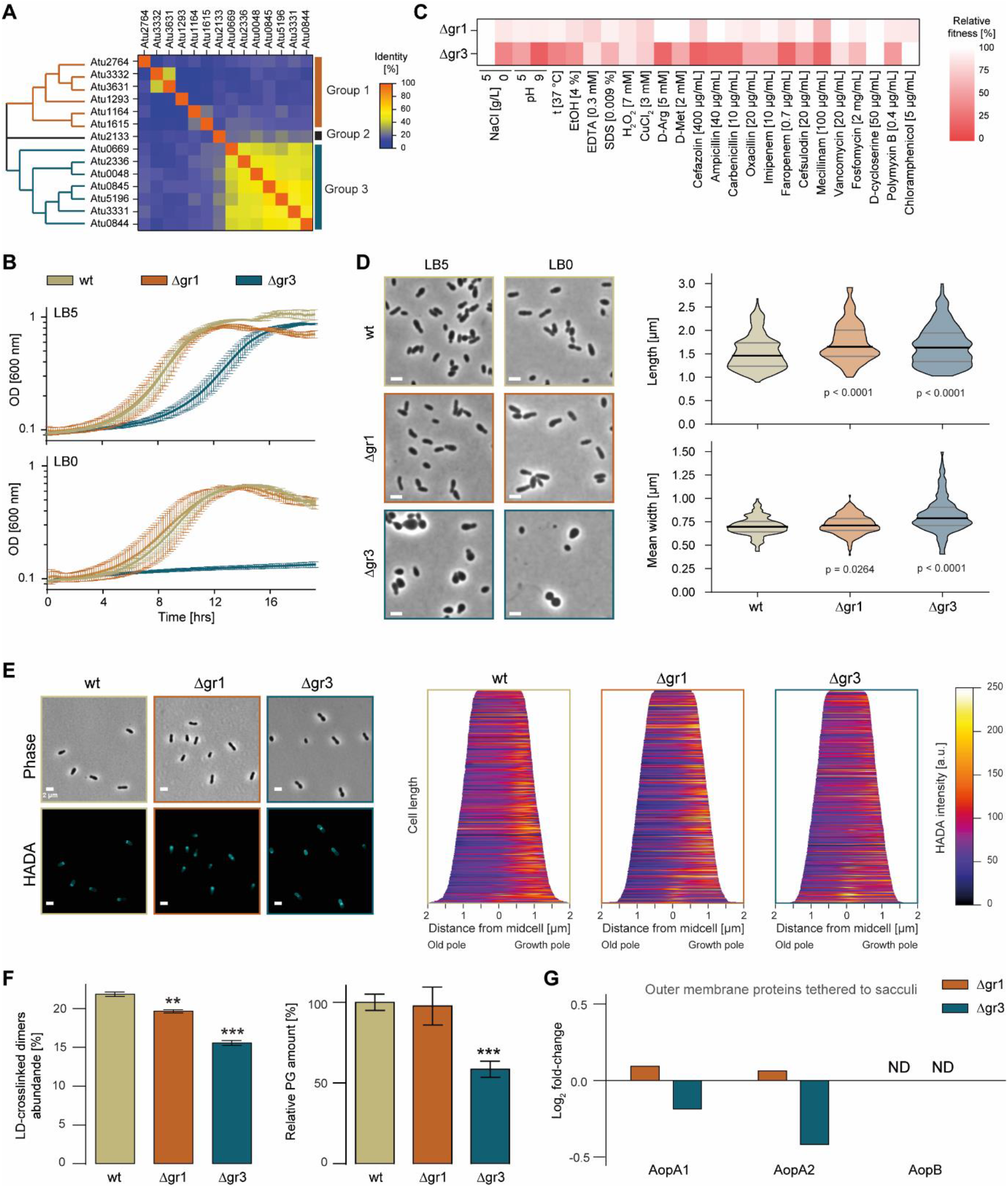
Hyphomicrobiales-specific LDTs are necessary for growth, shape maintenance and PG synthesis in *A. tumefaciens*. **(A)** Clustering and identity matrix for the 14 LDTs in *A. tumefaciens* identifies three groups. **(B)** Growth curves of *A. tumefaciens* wild type (wt), Δgr1 and Δgr3 mutants in LB5 (0.5% NaCl) and LB0 (0% NaCl) medium. **(C)** Relative fitness of *A. tumefaciens* Δgr1 and Δgr3 mutants (compared to wild type) under different conditions. **(D)** Representative phase contrast images of cultures grown in LB5 and LB0, and violin plots of the length and mean width of *A. tumefaciens* wt, Δgr1 and Δgr3 strains grown in LB5. Scale bar: 2 µm. **(E)** Representative phase contrast and fluorescence images of cultures (in LB5) treated with the fluorescent D-amino acid (FDAA) HCC-amino-D-alanine (HADA). Scale bar: 2 µm. Demographs depicting the incorporation of HADA at a population level are shown. **(F)** Relative abundance of LD-crosslinked dimers and relative PG amount in *A. tumefaciens* wt, Δgr1 and Δgr3 strains grown in LB5. Error bars represent standard deviation. **, p <0.01; ***, p <0.001. **(G)** Abundance of outer membrane β-barrel proteins (relative to wild type) known to be tethered to the PG in *A. tumefaciens* Δgr1 and Δgr3 strains grown in LB5.

In terms of morphology, Δgr3 cells appeared noticeably more spherical (wider and longer) compared to the rod-shaped wild type and the Δgr1 mutant in both media (Fig. 2D). In contrast to the more dispersed distribution of group 1 LDTs, several group 3 counterparts exhibit polar (Atu0048, Atu0844, Atu0845) or subpolar localization (Atu2336, Atu5196) (Fig. S4). Thus, we hypothesized that the morphological abnormalities observed in Δgr3 mutants might be related to reduced polar synthesis. To monitor PG synthesis, we tracked the incorporation of fluorescent D-amino acids (FDAA) (23, 24). Surprisingly, we observed no changes at the new pole, but detected an increased signal at the old pole (Fig. 2E). This suggests mislocalization and likely defective cell wall synthesis in the Δgr3 mutant. Indeed, while the abundance of LD crosslinked dimers decreased in both Δgr1 and Δgr3 mutants, the reduction was notably more pronounced in the Δgr3, with a decrease of 25% (Fig. 2F, S5). Furthermore, the relative amount of PG (normalized by optical density) decreased 50% in the Δgr3 mutant compared to wild type (Fig. 2F).

In addition to their role in forming PG crosslinks, LDTs can also catalyze the crosslinking of outer membrane β-barrel proteins (OMPs) to the PG (9). Therefore, considering that the observed PG defects likely contribute to the reduced viability and altered cell shape observed in the Δgr3 mutant, we investigated whether these mutants also exhibited impaired attachment of these proteins. To this end, we harvested sacculi and used a quantitative proteomics approach to determine the relative abundance of three OMPs known to be crosslinked to PG in this manner: AopA1, AopA2 and AopB (Atu1020, Atu1021 and Atu1131, respectively) (9). Although we were unable to detect AopB, the sacculi collected from the Δgr3 mutant had a lower abundance of both AopA1 and AopA2 compared to the wild type (Fig. 2G). We detected a small increase of both these OMPs in sacculi from the Δgr1 mutant. Interestingly, although strains with single or combined deletions of *aopA2* and *aopB* grew like the wild type strain and exhibited identical PG profiles (Fig. S6), constructing a Δ*aopA1* mutant was not possible, indicating this protein is likely essential.

In summary, our findings indicate that while both group 1 and 3 LDTs significantly contribute to maintaining LD-crosslinking in *A. tumefaciens*, only the Hyphomicrobiales-specific LDTs are indispensable for maintaining cell wall integrity and morphogenesis. These critical functions, which involve not only maintaining LD-crosslinking but also tethering the peptidoglycan to the outer membrane, cannot be sustained in the absence of group 3 LDTs from groups 1 and 2.

### Hyphomicrobiales specific LDTs are genetically linked to cell division factors

To reveal the genetic interactions of group 1 and 3 LDTs, we used a transposon insertion sequencing (Tn-seq) screen to assess each gene’s contribution to fitness in these mutant backgrounds. Few insertions resulted in conditional lethality or improved fitness in the Δgr1 mutant background (Fig. 3A, Supplementary File 2). Synthetic lethality included transposon insertions into the gene for inner membrane protein *cvpA,* the PG recycling ABC transporter *yej* (25), and *atu2682*, encoding for the Bax Inhibitor-1, a protein that has been associated with membrane homeostasis in the Alphaproteobacteria *Brucella suis* (26). Insertions that were conditionally beneficial in the Δgr1 strain included: the AMP nucleosidase *atu1006* and the putative acyl-CoA dehydrogenase *atu1310* genes. In the Δgr3 mutant, we observed a similar pattern of synthetically beneficial mutations as in the Δgr1 mutant. However, compared to Δgr1, the Δgr3 mutant displayed a broader range of synthetically essential genes. These genes included again *cvpA* and the *yej* transporter, but also several others related to cell wall biogenesis and division, such as the lytic transglycosylase *mltG*, the bifunctional PBPs *pbp1b1 and pbp1b2,* the DD-carboxypeptidase *dac* and *ftsK2* (Fig. 3BCD, Supplementary File 2). Consistently, Δgr3 mutant was found to be sensitive to the divisome inhibitor Aztreonam (Fig. 3E). Notably, insertions in the group 1 LDT *atu1164*, *E. coli*’s LdtD homolog, were also found to be synthetically lethal in the Δgr3 background, emphasizing the unique role of this LDT in preserving cell wall integrity when Hyphomicrobiales LDTs are absent. Additionally, the second most common COG term associated with under-represented insertions was for genes of unknown function (Fig. 3CD, COG category S). This suggests the presence of a pool of additional uncharacterized genes that are likely crucial for maintaining cell envelope biology.

**Fig. 3.**
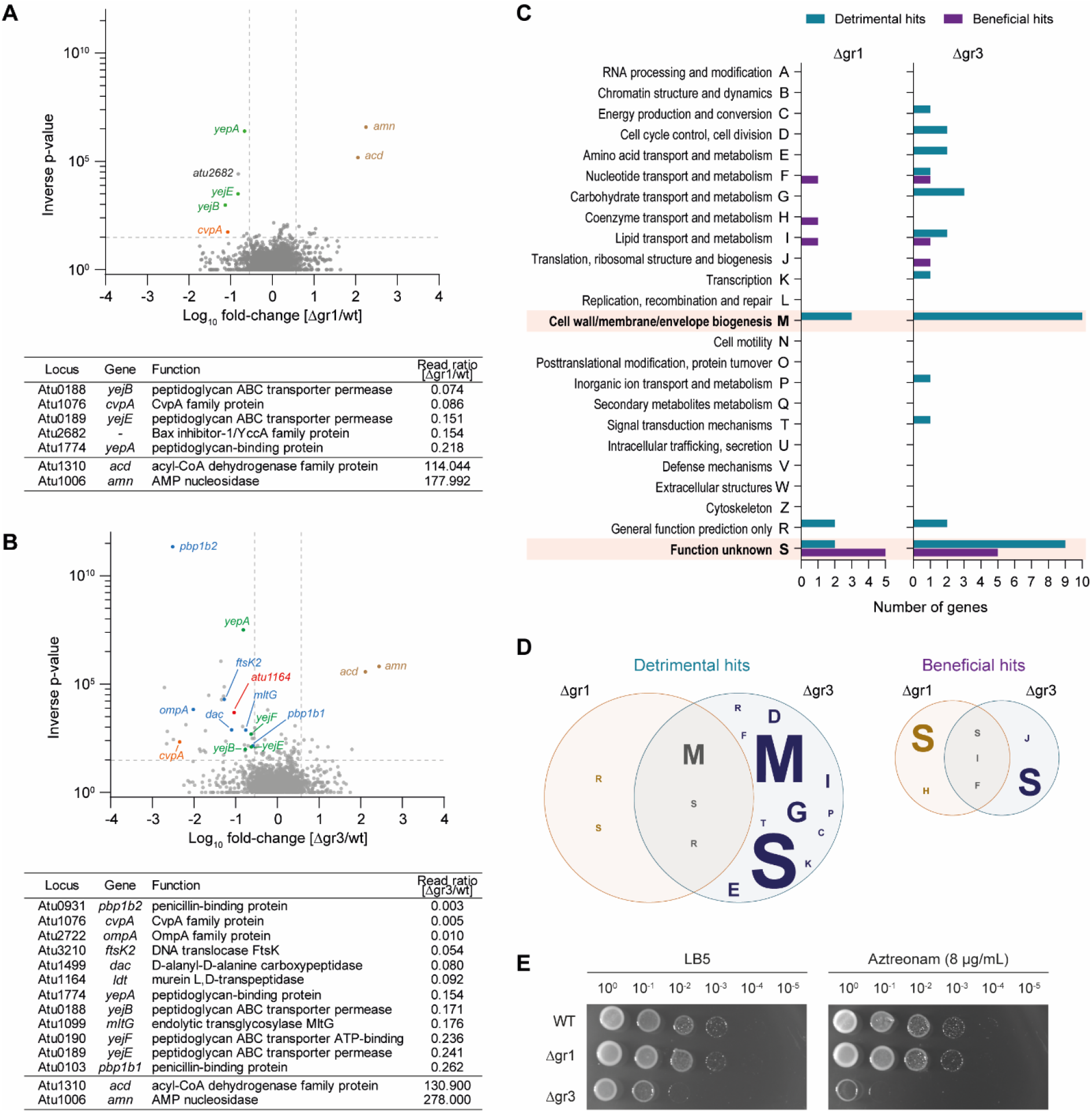
Genetic interactions of group 1 and group 3 LDTs. **(A)** Volcano plot showing the ratio of Transposon sequencing (Tn-seq) reads mapped to genes in the *A. tumefaciens* Δgr1 mutant strain and **(B)** Δgr3 mutant strain relative to wild type. Selected genes with synthetically detrimental (negative log-fold change) and synthetically beneficial (positive log-fold change) Tn insertions are highlighted. A table with the loci of interest is included. **(C)** Protein functions (COG functional classification) of the significantly synthetically detrimental and synthetically beneficial hits from Tn-seq experiments shown in A and B. **(D)** Venn diagrams representing the overlap of protein functions between the significantly synthetically detrimental (left) and synthetically beneficial (right) hits in the *A. tumefaciens* Δgr1 and Δgr3 mutant strains. The size of the letter is proportional to the number of genes within the specific COG functional classification. **(E)** Serial dilutions (10^0^ to 10^-5^) from overnight cultures of *A. tumefaciens* wt, Δgr1 and Δgr3 strains grown in LB5 spotted onto LB5 agar plates supplemented with Aztreonam 8 μg/mL. Growth on non-supplemented plate (LB5) was used as control.

Taken together, these results reinforce the idea that group 3 LDTs play a central role in peptidoglycan biogenesis, growth, and shape maintenance in *A. tumefaciens*. Additionally, they underscore the partial functional divergence of LDTs in this bacterium.

### Essentiality of LD-transpeptidation in *A. tumefaciens* is mediated by functionally diverse LDTs

The synthetic lethality observed between Δgr3 and Tn insertions intro the LDT gene *atu1164* underscored the heightened importance of group 3 LDTs and suggested that LD-transpeptidation might be essential in *A. tumefaciens*. In this light, certain LDTs appear more important for survival than others. To delve deeper into the functional redundancy of LDTs and ascertain the essential set of LDTs for *A. tumefaciens* viability, we attempted to engineer a strain devoid of all LDTs (referred to as Δ14). To minimize the risk of isolating suppressor mutations, we began combining Δgr1 and Δgr2, as these strains grew similar to the wild type. Subsequently, we deleted group 3 LDT genes sequentially. However, we could only generate a Δ13 mutant, which, based on the order of deletion, left only the group 3 LDT Atu3331 intact, leading us to designate this strain as Δ13 (Atu3331). This result indicates that LD-transpeptidation is vital in *A. tumefaciens*. Surprisingly, the phenotypes of Δ13 (Atu3331) were not exacerbated compared to those of the Δgr3 mutant, despite lacking several additional LDTs (compare Fig. 2 and Fig. 4ABC). Interestingly, MS-based quantitative proteomics showed that the abundance of Atu3331 was increased 2-fold in the Δ13 (Atu3331) mutant compared to wild type (Fig. 4D, Supplementary File 3). These results indicate that under LD-crosslink deficit, *A. tumefaciens* can boost the expression of remaining LDTs to maintain homeostasis of LD-crosslinking.

**Fig. 4.**
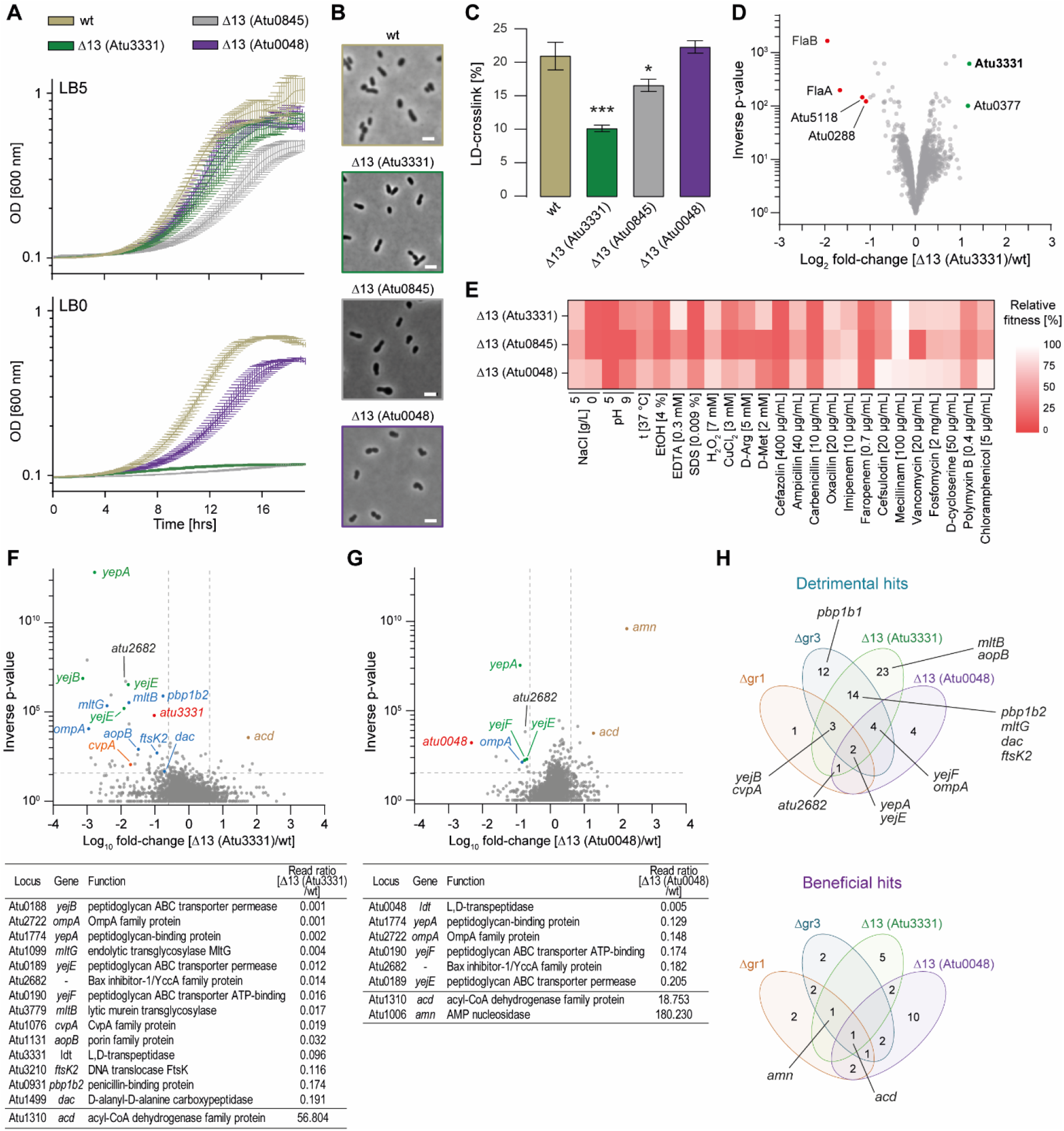
Mutants with a single LD transpeptidase can maintain fitness. **(A)** Growth curves of *A. tumefaciens* wild type (wt) and Δ13 mutant strains in LB5 (0.5% NaCl) and LB0 (0% NaCl) medium. **(B)** Representative phase contrast images of *A. tumefaciens* Δ13 *ldts* mutant strains grown in LB5. Scale bar: 2 µm. **(C)** Relative abundance of LD-crosslinked dimers and relative PG amount in *A. tumefaciens* wt and Δ13 *ldts* mutant strains grown in LB5. Error bars represent standard deviation. *, p <0.05; ***, p <0.001. **(D)** Volcano plot depicting the ratio of protein abundance of the *A. tumefaciens* Δ13 (Atu3331) mutant relative to wild type. Proteins shown in red and green have significantly lower and higher abundance, respectively. **(E)** Relative fitness of *A. tumefaciens* Δ13 *ldts* mutant strains (compared to wild type) under different conditions. **(F)** Volcano plot showing the ratio of Transposon sequencing (Tn-Seq) reads mapped to genes in the *A. tumefaciens* Δ13 (Atu3331) mutant strain and **(G)** Δ13 (Atu0048) mutant strain relative to wild type. Selected synthetically detrimental and synthetically beneficial hits are highlighted. A table with the loci of interest is included. **(H)** Venn diagrams representing the overlap of synthetically detrimental (top) and synthetically beneficial (bottom) hits in the *A. tumefaciens* Δgr1, Δgr3, Δ13 (Atu3331) and Δ13 (Atu0048) mutant strains.

To ascertain whether other LDTs alone could maintain *A. tumefaciens* viability in the absence of their homologs, we created two additional Δ13 mutants with different group 3 LDTs remaining: specifically, Δ13 (Atu0048) and Δ13 (Atu0845). While these alternative Δ13 mutants were also viable, our initial growth screening indicated important phenotypic differences, including variability in their ability to grow in standard and osmotically challenging conditions (Fig. 4A). Consistent with the growth data, the morphology and LD-crosslinking levels of the Δ13 (Atu0048) mutant were similar to those of the wild type strain while Δ13 (Atu0045), and particularly Δ13 (Atu3331), were more affected (Fig. 4BC, S7). To further investigate the function and degree of redundancy of the three terminal LDTs, we compared the growth phenotypes of these Δ13 mutants to the wild type strain under the same panel of sub-optimal growth conditions used previously (Fig. 4E, Supplementary File 1). All mutants were more sensitive to acidic pH, cefazolin, ampicillin, carbenicillin, faropenem and chloramphenicol. Individually, Δ13 (Atu0845) was more sensitive to vancomycin and pH 9 compared to the others, while the growth of Δ13 (Atu0048) was less affected than the other mutants across many of the conditions, including for example low salt, CuCl_2_ and cefsulodin. These results strengthen our previous conclusions about LDT activities of *A. tumefaciens* being partially specialized and further demonstrate a major role for Atu0048 in maintaining the cell wall integrity of this bacterium.

To evaluate the essentiality of the final LDT, we used Tn-seq in the Δ13 (Atu0048) and Δ13 (Atu3331) mutants (Fig. 4FGH, S8, Supplementary File 2). As expected, our results showed that insertions in the locus of the remaining LDT were highly under-represented in their respective Δ13 backgrounds, thus confirming that deleting all LDTs is lethal in *A. tumefaciens*. Notably, the two mutants exhibited important differences: the strain Δ13 (Atu3331) showed more synthetic interactions than Δ13 (Atu0048) and exhibited a synthetic lethality pattern that resembled that previously observed for the Δgr3 mutant (Fig. 4H). These hits included the PG recycling transporter *yejBEFyepA*, *pbp1b1 and pbp1b2*, *mltB* and *mltG*, *dac*, *ftsK2*, *aopB*, *cvpA* and *atu2682* for Δ13 (Atu3331) and *yejFEyepA*, *atu2682*, and *ompA* for Δ13 (*Atu0048*) (Fig. 4FG). These results further highlight the dominant role of Atu0048 among *A. tumefaciens* LDTs in cell wall homeostasis, fitness, and morphogenesis.

## Discussion

LDTs are present in both Gram-negative and -positive bacteria (11, 27–29); however, their primary function is considered ancillary. In *E. coli*, LDTs are non-essential, yet they enhance resistance to broad-spectrum β-lactams and reinforce the bacterial cell envelope in response to outer membrane defects (15, 16). Certain species, such as the Actinomycetales and Hyphomicrobiales, exhibit PGs with elevated LD-crosslinking. These species also encode LDTs that play crucial roles in growth and cell shape maintenance (17, 28, 30, 31). Notably, in *M. smegmatis*, the deletion of all LDTs results in a loss of rod shape in a subpopulation of cells and localized spherical blebbing due to a defect in cell wall rigidity (30).

In contrast to Actinomycetales, Hyphomicrobiales grow only from a single pole (17, 23, 32), and encode a high number of putative LDT proteins, with up to 20 predicted LDTs in some species. Also, it is common in members of this group, such as *A. tumefaciens*, to lack canonical elongation factors e.g., MreBCD, RodA, RodZ, and PBP2 (21). Previous studies propose that repurposed cell division components and LDTs could collaborate to promote polar growth in this bacterium. Specifically, canonical cell division proteins FtsZ and FtsA transiently accumulate at the growth pole, while the Hyphomicrobiales-specific LDT Atu0845 exhibits polar localization that correlates with FtsZ activity (21, 33, 34), thus supporting this hypothesis.

In our study, we found that *A. tumefaciens* requires at least one LDT for viability. Deleting the Hyphomicrobiales-specific LDTs significantly reduces LD-crosslinking, leading to severe elongation defects and cell rounding. While reduced LD-crosslinking could explain the phenotypes of the Δgr3 mutant, it is important to note that some LDT homologues also have distinct functions. For instance, among the 6 LDTs in *E. coli*, LdtD and LdtE specifically form LD-crosslinks (11), while LdtA, LdtB and LdtC are responsible for tethering the outer membrane-anchored Braun’s lipoprotein (Lpp) to the PG (10). Additionally, LdtF functions as an amidase, cleaving off Braun’s Lpp from PG (35). Recently, it was shown that although *A. tumefaciens* and other related species do not possess Lpp, they do tether their PG to various outer membrane β-barrel proteins (9, 36). In *Coxiella burnetii* and *Brucella abortus* this crosslinking is catalysed by LDTs (9, 36). Our results in *A. tumefaciens* suggest that this activity is catalysed mainly by Hyphomicrobiales-specific LDTs, as the Δgr3 mutant exhibits a significant reduction of PG-linked β-barrel proteins.

Our genetic screening revealed that deletion of Hyphomicrobiales-specific LDTs is synthetically lethal with the inactivation of *divK* and *ftsK2*, specific components of the coordination of division and development (CDD) pathway. While some CDD genes (e.g., CtrA) are essential, DivK is not. Deletion of *divK* disrupts FtsZ2 localization, resulting in branched and elongated rod-shaped cells in *A. tumefaciens* (37). Interestingly, transcription from CtrA activated promoters such as *ccrM* is increased in *divK* mutants (38). Since CtrA binding sites have not been identified in *A. tumefaciens*, we used the consensus binding sequences of CtrA from closely-related species to identify potential occurrences of the motif upstream of genes across the genome in silico (39–41). Remarkably, the CtrA binding motif is present upstream of group 1 LDT genes *atu1164, atu1293, atu1615, atu3332* and *atu3631*, in group 2 LDT *atu2133,* and in group 3 *atu0048, atu0669* and *atu2336,* suggesting that expression of some LDTs might be particularly relevant when regulation of the cell cycle is perturbed (Fig. S9).

Another intriguing result emerging from the Tn-seq screen was the synthetic lethality between the group 1 LDT Atu1164 in the group 3 LDTs. This finding suggests that Atu1164 uniquely contributes to viability when Hyphomicrobiales-specific LDTs are absent. One possible explanation is that certain LDTs are induced to help bacteria cope with stress, such as a weakening of the cell wall due to decreased crosslinking levels. In this line, we observed that *A. tumefaciens* increased the expression of Atu3331 when all the other 13 LDTs were inactive (i.e., in the Δ13 (Atu3331) mutant). This observation supports the idea that specific LDTs can be induced under stress conditions to compensate for LD-crosslinking defects. Notably, the ChvG/I pathway in *A. tumefaciens* modulates several LDTs and OM proteins. (42, 43). Similarly, expression of certain LDTs in other species is controlled by the general or cell envelope stress responses (8, 15). Although varying expression levels can enhance fitness to some extent, the phenotypic differences observed among three distinct Δ13 LDT mutants—each with a different LDT remaining as the last one—highlight the non-redundant functions of these proteins. It remains to be seen whether LDTs other than those belonging to the Hyphomicrobiales-specific group, particularly those that perform LD-crosslinks but do not tether PG to outer membrane β-barrel proteins, can support cell wall housekeeping functions in *A. tumefaciens*. Future investigations will focus on dissecting the specific contributions of each LDT to cell envelope integrity, conditional fitness, morphogenesis, and polar growth in *A. tumefaciens*. Advancing our understanding of the functional specialization of LDTs may yield valuable insights into other polarly growing Hyphomicrobiales species beyond *A. tumefaciens*, including human pathogens such as *Brucella abortus*. These insights could pave the way for novel antibacterial strategies that leverage the unique relevance of these enzymes in the biology of these bacteria.

## Materials and methods

### Media and bacterial growth conditions

Bacterial strains are listed in Supplementary Table S1.

Bacteria were routinely grown in Luria Bertani (LB) broth and agar plates (1.5 % (w/v) agar). When required, antibiotics were added to the culture medium or plates at the following concentrations: kanamycin 300 μg/mL for *A. tumefaciens*, and 50 μg/mL for *E. coli*. *A. tumefaciens* strains were grown at 30 °C, unless otherwise specified. *E. coli* strains were grown at 37 °C.

For growth curves, 100 µl bacterial cultures were grown in triplicate in 96 well plates at 30 °C (unless otherwise indicated), with orbital shaking. Optical density (absorbance at 600 nm, OD_600_) was measured every 10 minutes for up to 24 hours in a BioTek Eon Microplate spectrophotometer (BioTek, Winooski, VT, USA). For phenotypic growth screens, growth curves in LB medium supplemented with different compounds and conditions (Supplementary Table S5) were monitored. The relative fitness (%) of the wild type in each condition is calculated as the final maximal OD relative to that in standard lab condition (LB 0.5 % NaCl, pH7, 30 °C). Relative fitness of the mutants is calculated as the final maximal OD relative of the mutant strain relative to the wild type growth in each condition. Data is presented in Supplementary File 1.

Viability drop assays were done with normalized overnight cultures subjected to serial 10-fold dilution. Five-microliter drops of the dilutions were spotted onto control and Aztreonam 8 μg/mL supplemented LB agar plates and incubated at 30 °C for 24-48 h prior to image acquisition.

### Construction of mutants

Plasmids and primers are listed in Supplementary Tables S2 and S3 respectively.

For deletion of LDTs the upstream and downstream regions of the gene (about 500-600 bp) were amplified from purified genomic DNA with corresponding gene primers P1 and P2; P3 and P4 respectively. The upstream and downstream fragments were combined with corresponding P1 and P4 primers and inserted in pNPTS139 (44). The resulting plasmid pNPTS139*-*(*ldt* gene number) was confirmed by Sanger sequencing. In-frame deletion was introduced by allele replacement via homologous recombination (45). In short, exconjugants were obtained by conjugation and selected on ATGN plates with kanamycin 300 μg/mL and then subjected to sucrose counter-selection (46).

### Construction of LDT-sfGFP fusions

To construct expression vectors containing LDT-sfGFP, the respective coding sequence was amplified from purified genomic DNA. The amplicons were digested overnight and ligated into cut pSRKKM-Pcym using NEB T4 DNA ligase at 4 °C overnight, to create an expression vector compatible with the depletion strains (47). All expression vectors were verified by Sanger sequencing. All vectors were introduced into *A. tumefaciens* strains utilizing standard electroporation protocols (45).

### Protein structure analyses

Domain architecture was analyzed using Interpro (48).

Signal peptide predictions were performed with SignalP 6.0 (49).

Structural predictions were obtained from AlphaFold DB, version 2022-11-01 (50, 51). UCSF ChimeraX version 1.7.1 was used for visualization (52). Regions with pLDDT (predicted local distance difference test) lower than 50 have been hidden in the shown models.

### Analysis of LDT homologues

Unless otherwise specified, all software options were left default.

In order to identify LDT putative orthologues, we ran Orthofinder (version 2.4.0) (53) with non-default options [-M msa -A muscle -T iqtree]. The software defines “orthogroups” as genes which share and evolutionary origin. IQTree (version 1.6.12 multicore) (54) was configured to run with non-default options [-nt AUTO -safe -bb 1000 -bnni]. MUSCLE (version 3.8.31) was left default (55). Supplementary Table S4 contains the taxa, assembly versions and other meta data for all protein data files that were used in the analysis. Next, Interproscan (version 5.46-81) was run for all proteomes in Supplementary Table S4 with default parameters using InterPro release 81 (August 2020) [--iprlookup] (56). Then, we retrieved hidden markov model (HMM) profiles for YkuD (PF13645) and Ykud2 (PF03734) from the protein family database Pfam (release 33.1) (57). Both models were combined into a single input file and HMMER (version 3.3.1) (58) was run with non-default parameters [--noali]. All results from the Orthofinder, InterPro and HMMER analyses were then combined using a custom python script (Jupyter notebook and Python script: Supplementary Files 4 and 5), counting total number of LDTs per strain and matching orthologs of chosen strains *to A. tumefaciens* LDTs using Python version 3.8.

LDTs were clustered based on their protein sequence similarity by Multiple Sequence Comparison by Log-Expectation (MUSCLE) (55, 59). Identity matrix was built using the percentage of identity from the multisequence alignment.

### CtrA-binding motif analysis

CtrA-binding motifs TTAA-N7-TTAA and TTAACCAT (39–41) were used in FIMO (find individual motif occurrences) (60) with the 250 bp upstream of the *A. tumefaciens* LDT gene promoters. A P-value cutoff of 0.05 was used to establish the *A. tumefaciens* genes containing the motifs. Sequence logos were generated in R v4.3 using the ggseqlogo package (61).

### PG analysis

PG isolation and analysis were done as previously described (62–64).

Bacterial cells were pelleted by centrifugation (4,000 rpm, 20 min) and boiled in SDS 5% (w/v) for 2 h. Peptidoglycan was obtained by centrifuging for 13 min at 60,000 rpm at 20 °C (TLA100.3 Beckman rotor; OptimaTM Max ultracentrifuge Beckman, Beckman Coulter, California, USA). Pellets were washed 3-4 times by repeated cycles of centrifugation and resuspension in water. The washed pellet was digested with muramidase (Cellosyl 100 μg/mL) for 16 h at 37 °C. Muramidase digestion was heat inactivated and coagulated protein was removed by centrifugation for 15 min at 15,000 rpm. For sample reduction, pH of the samples was first adjusted to pH 8.5–9.0 with borate buffer, and then a freshly prepared NaBH_4_ 2 M solution was added to a final concentration of 10 mg/mL. After 20 min at room temperature, pH of the samples was adjusted to pH 3.5 with phosphoric acid and filtered (0.2 μm pore size filters).

Analysis of muropeptides was performed on an ACQUITY Ultra Performance Liquid Chromatography (UPLC) BEH C18 column, 130Å, 1.7 μm, 2.1 mm x 150 mm (Waters Corporation, USA) and detected at Abs. 204 nm with ACQUITY UPLC UV-visible detector. Muropeptides were separated at 45 °C using a linear gradient from solvent A (formic acid 0.1% (v/v) in water) to solvent B (formic acid 0.1% (v/v) in acetonitrile) in an 18 minutes run with a 0.25 mL/min flow.

Identity of the muropeptides was confirmed by MS and MS/MS analysis, using a Xevo G2-XS Q-tof system (Waters Corporation, USA). The QTOF-MS instrument was operated in positive ion mode. Detection of muropeptides was performed by MS^E^ to allow for the acquisition of precursor and product ion data simultaneously, using the following parameters: capillary voltage at 3.0 kV, source temperature 120 °C, desolvation temperature 350 °C, sample cone voltage 40 V, cone gas flow 100 L/h, desolvation gas flow 500 L/h and collision energy (CE): low CE: 6 eV and high CE ramp: 15-40 eV. Mass spectra were acquired at a speed of 0.25 s/scan over a range of *m/z* 100–2000. Data acquisition and processing was performed using UNIFI 1.8.1 software (Waters Corp.).

Chromatograms shown are representative of three biological replicates. Relative abundance of individual muropeptides was quantified from the relative area of the corresponding peak compared to the total area of the chromatogram. Unpaired t-test was used to statistically compare muropeptides’ abundance.

### Microscopy

Stationary phase bacteria were immobilized on 1% (w/v) agarose LB pads. Phase contrast microscopy was performed using a Zeiss Axio Imager Z2 microscope (Zeiss, Oberkochen, Germany) equipped with a Plan-Apochromat 63X phase contrast objective lens and an ORCA-Flash 4.0 LT digital CMOS camera (Hamamatsu Photonics, Shizuoka, Japan), using the Zeiss Zen 2 Blue software [v2.0.0.0]. Measurement of cells length and width was done in Fiji/ImageJ using MicrobeJ plug-in (65, 66).

For protein localization assays, cells containing plasmids with fluorescent protein fusions were grown at 28 °C in ATGN to exponential phase before imaging on agarose pads. When necessary, expression of plasmid encoded Ldt-sfGFP was induced by the presence of 0.2 mM cumate or 1 mM IPTG for 2 hours prior to imaging. A small volume (∼1 μl) of cells in exponential phase (OD_600_ = 0.4 to 0.6) was applied to a 1% ATGN agarose pad as described previously (67). Phase-contrast and epifluorescence microscopy was performed on ∼1000 cells across three biological replicates and representative images shown in Fig. S4.

For incorporation of FDAA, cells grown overnight at 28 °C in LB medium were diluted to an OD_600_ of 0.2 and allowed to grow until reaching an OD_600_ of 0.4 to 0.6. Cells were then labelled with 1 mM fluorescent D-amino acid (FDAA) HCC-amino-D-alanine (HADA) (23) for 5 minutes. Next, cells were fixed with ethanol to prevent further growth and washed with phosphate buffered saline (PBS). Phase-contrast and epifluorescence microscopy was performed on ∼1000 cells (887 WT cells, 1152 Δgr1, 853 Δgr3 cells) across two biological replicates and representative images shown in Fig. 2. Demographs were constructed using MicrobeJ (65). For demographs, cells were arranged from top to bottom according to their cell lengths, and each cell was oriented such that the new pole (defined as the cell pole with the higher fluorescence intensity as determined by HADA incorporation) was oriented to the right. The scale bar for the demographs represents intensity and ranges from 0 to 250 arbitrary units (a.u.).

### Transposon insertion sequencing

For the identification of synthetically lethal genes in selected *A. tumefaciens* genetic backgrounds, transposon insertion sequencing (Tn-seq) was performed as described previously (68). In brief, *A. tumefaciens* 9×10^4^ – 1×10^5^ transposon mutants were generated for each biological replicate of triplicates for wild type, Δgr1, Δgr3, Δ13 (Atu3331) and Δ13 (Atu0048) strains by conjugation of *A. tumefaciens* with *E. coli* SM10 λPIR carrying the transposon donor plasmid pSC189 (69). Mutant libraries were selected on LB plates containing kanamycin 500 µg/mL and streptomycin 25 µg/mL and pooled genomic DNA fragments were analysed using a MiSeq sequencer (Illumina, San Diego, CA, USA).

Insertion sites were identified, and statistical representation of transposon insertions was determined using the ConArtist pipeline (70). Synthetically detrimental and beneficial hits are listed in the Supplementary File 2.

### Proteomic analysis

For protein abundance measurements, cells were grown until stationary phase in triplicates, washed with PBS buffer once at 4 °C, and then pelleted at 3000 rpm for 8 min at 4 °C. Pellets were then resuspended in lysis buffer (2% SDS, 250 U/mL benzonase, and 1 mM MgCl_2_ in PBS) and boiled for 10 min at 99 °C. Samples were digested using a modified sp3 protocol (71, 72), and peptides were labelled with TMTpro (Thermo Fisher Scientific) as previously described (73). After pooling the samples together, they were fractionated to six fractions with high pH fractionation and injected on an Orbitrap Q-Exactive Plus (Thermo Fisher Scientific) coupled to liquid chromatography. Details on the run conditions and instrument parameters are described in (74, 75).

Mass spectrometry raw data was searched against the *Agrobacterium fabrum* FASTA file (UP000000813 downloaded from UniProt) using the Mascot 2.4 (Matrix Science) search engine and isobarquant (76). Protein abundance changes were determined using limma (77) by comparing mutant samples with wild type controls.

For quantification of the relative abundance of the three OMPs known to be crosslinked to PG, AopA1, AopA2 and AopB (Atu1020, Atu1021 and Atu1311, respectively), quantitative proteomic analyses were performed on Δgr1 and Δgr3 mutant strains and compared to wild type samples.

Changes in the proteome of mutant strain Δ13 (Atu3331) are shown in Fig 4D and data is provided in Supplementary File 3.

### Statistical analysis

All statistical analyses were performed using GraphPad Prism (GraphPad Software, San Diego, CA, US). Student’s unpaired *t* tests (unpaired, two-tailed) were used to assess statistical significance. Assays were performed with three biological replicates unless otherwise indicated.

## Acknowledgements

We thank all the members of the Cava lab for helpful discussions and Dr. Amelia Randich for feedback on this manuscript. **Funding:** Research in the Cava lab is supported by The Swedish Research Council (VR), The Knut and Alice Wallenberg Foundation (KAW), The Laboratory of Molecular Infection Medicine Sweden (MIMS) and The Kempe Foundation. B.S. is supported by the Knut and Alice Wallenberg Foundation and the Vinnova UPSC Centre for Forest Biotechnology. Research in the Brown lab was supported by the National Science Foundation, IOS1557806, to P.J.B.B and University of Missouri Research Council Grant, URC-23-010. A.M. was supported by the Swedish Research Council/Vetenskapsrådet (grant number: 2022-02958), and Kempestiftelserna (grant number: JCK3126).

## Author contributions

Study conceptualization: A.A. and F.C. Experimental design: A.A., T.G., P.J.B.B and F.C. Performed the experiments: A.A., T.G., M.C.G., J.A., A.M. and I.R. Analysed the data: A.A., T.G., L.A., M.C.G., J.A, A.M. and B.S. Wrote the paper: A.A., T.G., L.A. and F.C. All authors commented on the manuscript. Supervision: A.T., M.M.S., P.J.B.B. and F.C. Funding acquisition: A.M., A.T., M.M.S., P.J.B.B. and F.C.

## Data and materials availability

All data needed to evaluate the conclusions in the paper are present in the paper and/or the Supplementary Materials. Additional data related to this paper may be requested from the authors.

